# Deciphering enzymatic potential in metagenomic reads through DNA language models

**DOI:** 10.1101/2024.12.10.627786

**Authors:** R Prabakaran, Y Bromberg

## Abstract

Microbial communities drive essential global processes, yet much of their functional potential remains unexplored. Metagenomics stands to elucidate this microbial “dark matter” by directly sequencing the microbial community DNA from environmental samples. However, the exploration of metagenomic sequences is mostly limited to establishing their similarity to curated reference sequences. A paradigm shift - language model (LM) -based methods - offer promising avenues for reference-free analysis of metagenomic reads. Here, we introduce two LMs, a pretrained foundation model REMME, aimed at understanding the DNA context of metagenomic reads, and the fine-tuned REBEAN for predicting the enzymatic potential encoded within the read-corresponding genes. By emphasizing function recognition over gene identification, REBEAN labels gene-encoded molecular functions of previously explored and new (orphan) sequences. Even though it was not trained to do so, REBEAN identifies the gene’s function-relevant parts. It thus expands enzymatic annotation of unassembled metagenomic reads. Here, we present novel enzymes discovered using our models, highlighting model impact on our understanding of microbial communities.

## Introduction

The world we live in is estimated to host up to a trillion bacterial species spread wide and deep across the globe (Locey & Lennon, 2016; Louca et al., 2019). Microbes are the foundation of Earth’s biosphere and are responsible for much of life as we know it. We learned of the existence of microbes with the discovery of the microscope in the 1600’s (Lane, 2015). Centuries later, however, we are only capable of isolating, growing, and studying a tiny fraction of these organisms in the lab (Ruscheweyh et al., 2022; Solden et al., 2016; Staley & Konopka, 1985; Steen et al., 2019).

An alternate to the exploration of individual microbes is studying them directly within their communities via the analysis of metagenomes, i.e. the bulk genetic material of environmental samples. Current applications of metagenomic studies are diverse, including but not limited to surveillance of antimicrobial resistance (AMR), exploring microbial adaptation to environmental changes, studying the influence of host microbiome on host health, identification of new microbial species, and discovery of novel genes and gene products (Handelsman, 2004; Ko et al., 2022; Zhu et al., 2018). Significant advances in sequencing techniques of the past two decades have largely overcome the technical challenges associated with metagenomic data extraction, shifting the focus towards downstream analyses, such as gene prediction, taxonomic classification, and functional annotation/profiling (Bharti & Grimm, 2021).

Functional profiling is an important step in deciphering the contents of the microbial sample (Ko et al., 2022; Pushkarev et al., 2018; Zhu et al., 2020). It provides a way to track molecular functionality known to be important and to identify novel genes of known or yet-undescribed functions. Functional labels used in profiling are predefined terms and are part of extant ontologies and domain/family collections, e.g. Enzyme Commission numbers (EC), Gene Ontology (GO) terms, Pfam protein families and InterPro signatures, Clusters of Orthologous Genes (COG), and KEGG genes and pathways (Ashburner et al., 2000; Kanehisa et al., 2022; Mistry et al., 2021; Paysan-Lafosse et al., 2023; Tatusov et al., 1997; Tipton, 1994). Functional annotation of a metagenomic sample may involve labelling whole genes, identified from assembled stretches of sequence (transcripts, contigs, or genomes), or inferring putative labels directly from sequencing reads (Bharti & Grimm, 2021).

Most current metagenome annotation methods rely on mapping of genes or reads to related reference, i.e. annotated, gene or product protein sequences. Reference-based function assignments use sequence alignment and k-mer indexing and matching (Aziz et al., 2008; Mistry et al., 2021; Overbeek et al., 2005; Sayers et al., 2024; Tatusov et al., 2000), Hidden Markov Models (HMMs) of families, or structural comparisons. Requiring an existing reference, however, limits the range of discovery possible with these methods (Prabakaran & Bromberg, 2025).

Recent developments in deep learning models, and particularly language models (LMs), have succeeded in predicting protein structure and made significant advances in protein domain and function annotations (Bileschi et al., 2022; John Jumper et al., 2021; Lin et al., 2023; Sanderson et al., 2023). These have been experimented with in the metagenomic world as well (Hoarfrost et al., 2022; Karin & Steinegger, 2025; Pan et al., 2022), suggesting that they can generalize the information encoded in sequencing reads. In this study, we trained REMME (Read Embedder for Metagenomic exploration), a foundational transformer-based DNA language model (dLM), to learn the “language” of sequencing reads. REMME is adaptable to various downstream tasks, and we describe one such task here: REBEAN (Read Embedding Based Enzyme Annotator) performs reference- and assembly-free annotation of enzymatic activities putatively encoded by microbial genes that give rise to metagenomic sequencing reads. REBEAN demonstrates robust performance, by leveraging understanding on the read context within their “parent” enzymes. Our analyses describe REBEAN’s extensive applicability to metagenome annotation, particularly highlighting its ability to forgo sequence-defined homology in favor of discovering novel enzymes.

We note that much of the recent emphasis in the protein and nucleotide work with dLMs has focused on creating novel sequences that perform desired functions (Madani et al., 2023; Nguyen et al., 2024). This approach, however, discards the well-optimized natural results of billions of years of evolution in favor of synthetic, often questionable, biology. Here we suggest that annotation of the existing rich diversity of sequences, rather than de novo synthesis, should be the first step in harnessing the power of biological molecules.

## Materials and Methods

### Model Development

In this work we introduced two DNA language models (dLMs): (i) a general-purpose, pretrained model that embeds DNA reads to capture underlying “language-like” patterns in nucleotide sequences, and (ii) a fine-tuned model for enzymatic function annotation, classifying reads into seven first level Enzyme Commission (EC) classes.

### REMME - Read Embedder for Metagenomic exploration

#### Data

We obtained 1,496 genomic assemblies representing 1,496 prokaryotic species from the marine microbiome samples in MGnify (Richardson et al., 2023). From these assemblies, we extracted 72.9 million reads (72,966,774) by randomly sampling 20 reads per 1Kbp with an average length of 136 bp (60-300 bps), as described in (Hoarfrost et al., 2022). The fraction of coding region residues in each read, along with the read position relative to the start codon were noted. Of these ∼73 million reads, 65.1M and 7.8M reads were from coding and non-coding region, respectively. We clustered this dataset at 80% sequence identity using MMseqs [v13.45111, (Hauser et al., 2016) ] to obtain 53.6M representative reads. These were further split into training (27.9M coding and 3.7M non-coding representatives of 40.2M coding and 4.7M non-coding), validation (12M coding and 1.6M non-coding representatives of 17.2M coding and 2M non-coding), and testing (7.3M coding and 1.1M non-coding representatives of 7.7M coding and 1.2M non-coding) sets, for an 59-25-16% split of representative sequences, respectively.

#### Training

REMME is an encoder-only transformer model, holding six transform layers with eight multi-attention heads. Each input DNA sequence was transformed into a sequence of overlapping nucleotide triplets (tokens) with a stride of one nucleotide. The tokens were fed into a token embedder that encodes each token in the sequence as a numeric vector of length 128 and a position embedder that encodes the sequence position of each token as a vector of length 128. The combined encoding from token embedder and position embedder was fed into the encoder module containing six encoder layers with eight multi-head attention heads. We had used GELU activation along with a drop out of 0.10 throughout the model everywhere, unless specified otherwise. The total number of trainable parameters was 1,662,177.

The encoder model was trained to perform masked-token prediction. Here, 15% of the nucleotides to the encoder input were perturbed, similar to BERT (Devlin, 2018) model training (80% masked and 10% random). The transformed read embeddings from the encoder module were then fed into three modules: 1) a *decoder* module, of three linear layers reconstructing the original nucleotide triplet sequence, 2) a *regression* module, comprising a 2d Fractional Max pooling layer, combined with two dense layers to predict the fraction of coding/non-coding residues in the read, and 3) a four-class *classifier* module, predicting the reading frame of the coding DNA sequence (CDS) that overlaps with a given read. The four classes represent three reading frames (classes 1, 2 & 3) and the non-transcribed reads (class 4). The losses from all three modules were summed and back propagated during training. The model was trained for 53 epochs until no significant decrease in total loss was observed.

### REBEAN - Read Embedding Based Enzyme Annotation

#### Data

We collected 306 studies containing 19,316 diverse metagenomic samples of non-viral origin from the recent update to the SRA database (20230627) (Katz et al., 2022). Among these, 3,136 metagenomic samples are from microbiomes associated with soil, water, coastal water, sea water, oligotrophic water, deep marine sediment, phyllosphere, algal, and anthropogenic material, we randomly selected 40 SRA experiment runs from each of the nine environments, yielding 360 runs representing 332 samples and containing 267.3M reads (**Table S2**).

The mi-faser method (Zhu et al., 2018), using 41,640 enzyme sequences as reference, was used to annotate these metagenomic reads with an enzymatic activity assigned to their putative ‘parent’ genes, i.e. gene sequences from which the read is taken. The enzyme reference set comprised genes (extracted from NCBI RefSeq (O’Leary et al., 2016)) encoding SwissProt proteins with Enzyme Commission (EC) number annotations and experimental evidence of protein existence (Sayers et al., 2024; UniProt, 2023). Note that only a third (14,229) of these enzymes had experimental evidence of their enzymatic activity. Mi-faser annotated 59.5M of 267.3M reads (22%) as belonging to enzymes. We extracted reads of length 60 to 300 bps and clustered them, at 80% sequence identity using MMseqs (v13.45111), to retain 16.6M (16,624,341) and 112.3M (112,334,039) sequence-dissimilar reads of enzymatic and non-enzymatic origin, respectively. To create a balanced class distribution for training, we randomly sampled 2.4 million non-enzymatic reads – a number representative of the average number of reads in each EC class. Final composition of the 19 million read dataset was 17.7% of oxidoreductase (EC 1), 25.9% of transferases (EC 2), 11.3% of hydrolases (EC 3), 8.8% of lyases (EC 4), 5.8% of isomerases (EC 5), 10.5% of ligases (EC 6), 7.6% of translocases (EC 7), 12.5% of non-enzymes. We held out 3.8M (20%) of the reads as a test dataset and split the rest of the data (15.2M) into training (13.7M, 90%) and validation (1.5M, 10%) without altering the proportion of each enzyme class.

#### Training

REBEAN is a version of REMME that was fine-tuned to label reads that come from ‘parent’ enzyme-coding genes. REBEAN consists of the six encoder layers from REMME coupled with one classifier module comprising three dense layers. The classifier annotates a given read as being class one through eight, representing seven first level EC classes and non-enzymes, annotated by mi-faser as described above. We trained REBEAN using an ADAMW optimizer and cosine restarts scheduler. The model was trained for 188 epochs until no significant loss decrease was observed. To mitigate bias arising from variable read lengths in metagenomic data, model training was restricted to the first 106 nucleotides of each read. This length threshold corresponds to the mean minus one standard deviation of the overall read length distribution.

### Model Evaluation

#### Performance measures

We used multiple standard metrics such as accuracy, recall, precision, specificity, F1 score, balanced accuracy, mean squared error (MSE) and mean absolute error (MAE) (**Eqn. 1-8**) to evaluate predictive performance of REMME and REBEAN in training and several independent analyses. We calculated the MSE (**Eqn. 2**) and the MAE (**Eqn. 3**) to quantify the difference between “ground truth” (f) and predicted coding fraction (f^p^) of reads. We also computed the Area Under the Curve (AUC, **Eqn. 9**) for Precision-recall (PR-AUC) and Receiver operating characteristic (ROC-AUC) curves (Fabian et al., 2011).

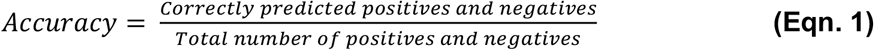

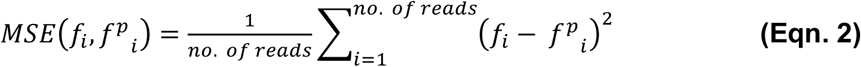

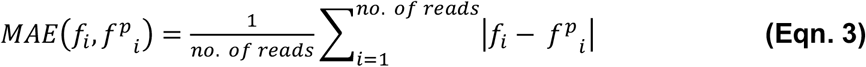

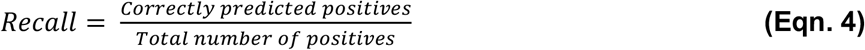

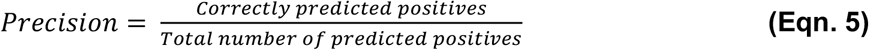

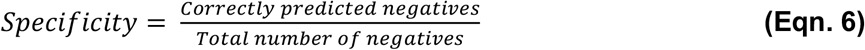

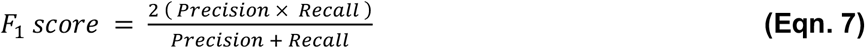

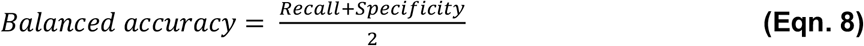

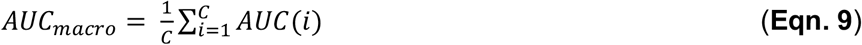

### Embeddings similarity between reads

We used cosine similarity and Euclidean similarity for comparing read embeddings (**Eqn. 10-11**). Apart from the inherent differences in similarity distributions, our inferences were consistent across these metrics. For clarity, we only show the results of Euclidean similarity unless specified otherwise.

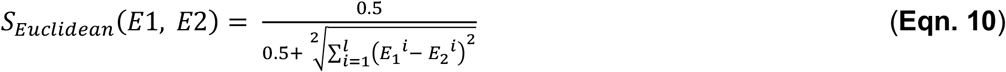

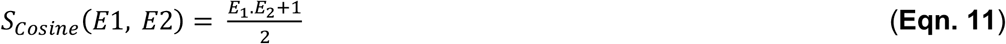

In several comparison, we computed embedding similarity of a read against a set of reads representing a gene, gene set, genome or metagenome. In these cases, we represent the multiple read-pair similarities as an aggregate score using mean and maximum statistics. For a read *k* compared against a set of *n* reads, the aggregate score is defined using **Eqn. 12 and 13. Eqn. 12** depicts the average embedding similarity of the read *k* against the read set representing a gene or genome. **Eqn. 13** represents the embedding similarity with the best-aligned read in the read set.

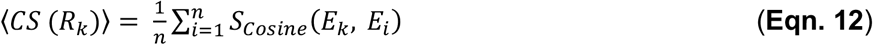

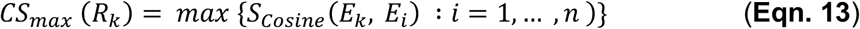

### Genes-to-synthetic reads

For multiple analysis, we generated synthetic reads from gene sequences to simulate metagenomic samples (**Table 1**). Unless otherwise specified, we randomly sampled, with no restrictions on overlap or coverage, ten gene fragments of length 200 nucleotides per 1Kbp of a gene. For example, for a gene of length 2Kbp, the number of reads sampled would be 20. This process of reads generation was also employed to generate reads from genomes.

**Table 1:** List of sequence datasets used in analysis.

| Dataset name | Description | Type | Number of entries | Number of reads | Synthetic reads<sup>1</sup> | Read sampling rate (reads/kbp) |
| --- | --- | --- | --- | --- | --- | --- |
| SPEnzset1 | Reads from SwissProt Exp. Enzymes I | Gene | 4,295 | 525,775 | Yes | 100 |
| SPEnzset2 | Reads from SwissProt Exp. Enzymes II | Gene | 4,295 | 54,115 | Yes | 10 |
| SPset | Reads from SwissProt Enzymes and Non-enzymes | Gene | 473,854 | 5,305,894 | Yes | 10 |
| Orthoset1 | Reads from Ortholog genes from OrthoDB | Gene | 24,938 | 707,947 | Yes | 10 |
| Orthoset2 | A subset of Orthoset1 | Gene | 7,311 | 79,738 | Yes | 10 |
| MarineMGset | Synthetic Metagenome from MGnify MAGs | Genomes | 3,820 | 124,269,382 | Yes | 10 |
| ExtremeMGset | Extremophile metagenomes | Metagenome | 25 | 8,035,798 | No | - |

<sup>1</sup>Synthetic reads were sampled from gene and genomes sequences as described in the section "Genes-to-synthetic reads"

### UniProt annotated enzyme read dataset

We validated the performance of REBEAN – the fine-tuned variant of REMME for first-level EC class prediction – using a curated set of experimentally validated enzyme sequences. From UniProt, we extracted 12,277 manually curated enzymes annotated with one unique EC number and experimental evidence of enzymatic activity (ECO:0000269) and mapped these to gene sequences using NCBI Entrez (Sayers et al., 2024; UniProt, 2023). Of these, we selected the 4,295 enzymes belonging to prokaryotes and archaea. These included 1,096 (EC 1), 1,300 (EC 2), 785 (EC 3), 509 (EC 4), 315 (EC 5), 216 (EC 6), and 74 (EC 7) enzyme sequences. We further generated a dataset of 525,775 reads by randomly sampling 100 reads of length 200 bp per 1Kbp of a gene (as described in section **Genes-to-synthetic reads)**. Each read was labelled with the 1^st^ level enzyme class of the parent gene. Among the half million reads, 129,111 (24.6%), 160,065 (30.5%), 95,638 (18.2%), 60,325 (11.5%), 35,476 (6.7%), 34,365 (6.5%) and 10,795 (2.1%) were annotated as EC class 1 to 7, respectively, based on the parent gene. This dataset of 545,775 reads (**SPEnzset1**) was used to access REBEAN’s prediction performance. The performance statistics were computed by sampling 10% of the reads over 100 independent iterations to simulate a read sample of 10 reads per 1Kbp. We also created additional read datasets by varying the sampling rate (1, 2, 5, 10, 20, 50 and 100 reads per 1Kbp) and read length (50, 100, 150, 200, 250, 30 and 400 bp) to assess the influence of sequencing depth and other parameters.

### SwissProt annotated enzyme versus unannotated proteins

To evaluate REBEAN’s ability to differentiate enzymes and non-enzymes, we collected gene sequences of 473,854 SwissProt proteins, as described above. Among these proteins, 231,203 were annotated as enzymes based on *in silico* or experimental evidence, while the remaining 242,651 were unannotated. From these 473k genes, we generated 5,305,894 synthetic reads of 200 bp in length (**SPset)**, including 2,965,689 reads derived from the 231,203 enzyme-annotated genes.

### Ortholog dataset

To further assess the representational ability of REBEAN and REMME, we analysed read-level embeddings derived from orthologous and non-orthologous gene pairs. We randomly selected 1,000 different ortholog groups (OGs) from OrthoDB [v11, (Kuznetsov et al., 2023)] at each of the five taxonomic levels: genus, family, order, class and phylum. For each OG, we randomly chose five genes and identified ten random ortholog gene pairs between them. We then formed ten non-ortholog pairs by pairing genes across different OGs of the same taxon. The final dataset comprised 65,169 unique genes which includes 24,938 genes in 50,000 orthologous gene pairs and 65,169 genes in 50,000 non-orthologous gene pairs. We mapped these 65,129 genes to EMBL CDS to obtain respective gene sequences. 707,947 reads (**Orthoset1)** were sampled from these 65,129 genes as described above (as described in section **Genes-to-synthetic reads)**.

Further, we selected a subset of 7,311 genes (79,783 reads, **Orthoset2**) from 100 ortholog groups for in-depth analysis of read pair embedding similarity. For the ∼6.4B corresponding read pairs, we computed sequence identity using Smith-Waterman (Cock et al., 2009; Smith & Waterman, 1981) with match, mismatch, gap scores 1, -2 and -1, respectively.

### Synthetic Metagenome Read dataset

To demonstrate the practical utility of REBEAN, we applied it to marine metagenome sequencing data to identify candidate oxidoreductases based on their read-level enzymatic signatures. We collected 3,820 marine metagenome-assembled genomes (MAGs) from MGnify (Gurbich et al., 2023) and generated 124 million genome fragments of 200bp length resembling metagenome reads (**MarineMGset)**. We sampled ten reads per 1Kbp of a genome without restrictions on overlap. In addition to the genome sequences, we retrieved the corresponding protein-coding sequences and associated annotations from MGnify (pipeline versions 4.0, 4.1, and 5.0). Functional and taxonomic annotations were derived using multiple tools integrated in the MGnify workflow, including EggNOG-mapper, HUMAnN, HMMER, and KEGG pathway mappings (Eddy, 2011; Gurbich et al., 2023; Huerta-Cepas et al., 2019; Richardson et al., 2023; Seemann, 2014).

Additionally, we ran HMMER [v3.4, (Eddy, 2011)] to assign Pfams to the putative oxidoreductases identified by REBEAN, with E-value threshold set at 1E-3. To assess the enrichment of oxidoreductase-associated Pfams in REBEAN-filtered proteins, we compared them to Pfams assigned to oxidoreductases in SwissProt (release 2023-11) (**Table S3**). Note Pfams of unknown function (DUFs) were excluded from this analysis. Enrichment was quantified using the odds ratio, and statistical significance was determined using the Hypergeometric test (*Categorical Data Analysis*, 2002; Virtanen et al., 2020).

### Metagenomes from extreme environments

To evaluate REBEAN’s performance in environments where organisms bear limited homology to reference databases, we collected 803 million metagenomic reads from 18 samples (25 Runs, **Table S4**) from three extremes environments (hydrothermal vents, hypersaline and salt crystallizer ponds) from MGnify. To speed up evaluation, a random 1% subsample of reads was selected from this dataset, excluding 301 reads that shared >30% identity (Hauser et al., 2016) with sequences in REBEAN’s training data, resulting in a final dataset of 8,035,798 reads (referred to as **ExtremeMGset**).

In addition to the sequencing data, coding sequences (CDSs) and corresponding Gene Ontology (GO) annotations were retrieved from MGnify (pipeline versions 4.0, 4.1, and 5.0) reflecting annotations of ∼500 million predicted protein sequences (**Table S4**). GO terms were mapped to Enzyme Commission (EC) numbers using the GO2EC database [GO2EC (Gene Ontology et al., 2023)].

We compared REBEAN’s predictions for this **ExtremeMGset** with those of existing pipelines: HUMAnN3 (using the UniRef90 release v201901b), mi-faser (v1.63), Carnelian, and LookingGlass (Beghini et al., 2021; Hoarfrost et al., 2022; Nazeen et al., 2020; Zhu et al., 2018). As HUMAnN3 outputs gene family-level functional profiles in Reads Per Kilobase (RPK) rather than direct read-level annotations, to estimate per-read EC counts, RPK values were multiplied by the corresponding gene family lengths. All tools were run using their default settings.

We further constructed an alignment-based baseline method by assigning reads to EC classes through protein sequence matches using DIAMOND (v2.1.10.164). Reads were aligned to UniRef90 (release 201901b) using a sequence identity cutoff of 90% and an alignment coverage cutoff of 50%. A total of 599,923 reads aligned to UniRef90, of which 155,573 matched proteins annotated as enzymes. For the 2,910 reads aligning to multiple enzyme classes, a single EC class was assigned based on the frequency of matches and alignment quality, prioritizing identity, bit score, and alignment length. This DIAMOND-based annotation provides a realistic upper bound for coverage achievable using traditional homology-based methods. EC annotations for UniRef90 entries were obtained from the HUMAnN3 reference database.

To account for updates in Enzyme Commission (EC) classifications, we revised the EC annotations according to the definitions provided by ExPASy (release 2019_07) (Bairoch, 2000; Tipton, 1994). For example, we relabelled two reads originally annotated by Carnelian as EC 1.6.5.8 to the updated classification EC 7.2.1.1. As our benchmarking is limited to first-level EC classes, these updates had minimal impact on the annotation results across the 8 million reads. The most notable change was the introduction of Class 7 (Translocases) in 2018.

### Assessment of method agreement

The agreement between two tools in annotating each of the 8M reads (**ExtremeMGset)** was evaluated by measuring the Cohen’s Kappa score **(Eqn 14)**. *P*_*e*_ and *P*_*o*_ represent the expected and observed proportion of agreement derived from marginal probabilities. Kappa values range from -1 (perfect disagreement) to +1 (perfect agreement).

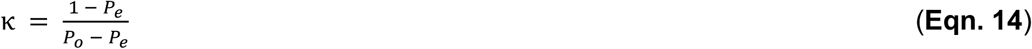

### Embedding visualization

We used TSNE for dimensionality reduction of the embedding space. TSNE 2D-projections were computed using (Policar et al., 2024) with following parameters: n_neighbors = 15, min_dist = 0.1 using *cosine* metric.

### Statistical Analysis

Statistical significances were assessed through the Wilcoxon rank-sum test and Student’s t-test using SciPy (Virtanen et al., 2020). Where applicable, 95% confidence intervals (CIs) for mean and median values were estimated via bootstrapping, using 90% subsamples of the data without replacement over 100 iterations.

## Results and Discussion

Here, we developed a transformer-based language model to annotate enzymatic activity of genes giving rise to sequenced reads. That is, given a particular read, we asked: “What would be the function of the gene that it came from?”

### Building a DNA-language model (dLM) to decipher metagenomic reads

Our first (foundation) model was pretrained as a generalized read embedder to understand the DNA “language” using 53.6 million reads from 1,496 prokaryotic genomes (**Methods**). It was then fine-tuned on 19 million non-redundant reads from 19,316 metagenomic samples to annotate enzymatic function in terms of the seven first level EC classes. For clarity of the discussion, we named the pretrained dLM REMME (Read Embedder for Metagenomic exploration) and the fine-tuned model REBEAN (Read Embedding Based Enzyme Annotator).

REMME was trained to embed DNA reads through mask-token prediction, i.e. predicting masked nucleotides in each read. It attained over 98.5% accuracy (**Eq. 1**) in each: training, validation, and testing (**Figure S1**). This performance highlights REMME’s solid understanding of the DNA “spelling”, i.e. the nucleotide order of occurrence. The nature of read data, i.e. short genomic fragments that lack in biological context, however, raised a concern that this model learned shallow sequence patterns, such as nucleotide composition and base frequencies, rather than meaningful biological features. To enforce further understanding of biological context, we gave REMME two additional learning tasks: (1) estimate the fraction of protein-coding nucleotides in each read and (2) if the read is part of a coding DNA sequence (CDS), predict the open reading frame; as part of the latter task, REMME was trained to predict a given read class: 1, 2, or 3 for each of the three-reading frames (with respect to the first position of the read in the 5’ to 3’ direction) and 4 if the read was not transcribed.

REMME learned to identify coding regions within reads well. In both validation and testing, the REMME-predicted fraction of coding nucleotides per read was well correlated with the ground truth fraction of coding nucleotides (**Figure 1A;** Pearson’s r = 0.73, *p-value* < 1E-32; MSE=0.04 and MAE=0.11, **Eqn. 2-3**). Note that the largest number of mispredicted coding fractions was attributed to mostly non-coding reads, i.e. those containing 0-0.1 known coding residues. This observation is in line with the known difficulty of distinguishing short coding and non-coding stretches (Clamp et al., 2007; Zhang et al., 2016). As the ground truth fraction of coding residues increased, REMME was able to identify the stretches more precisely.

**Figure 1:**
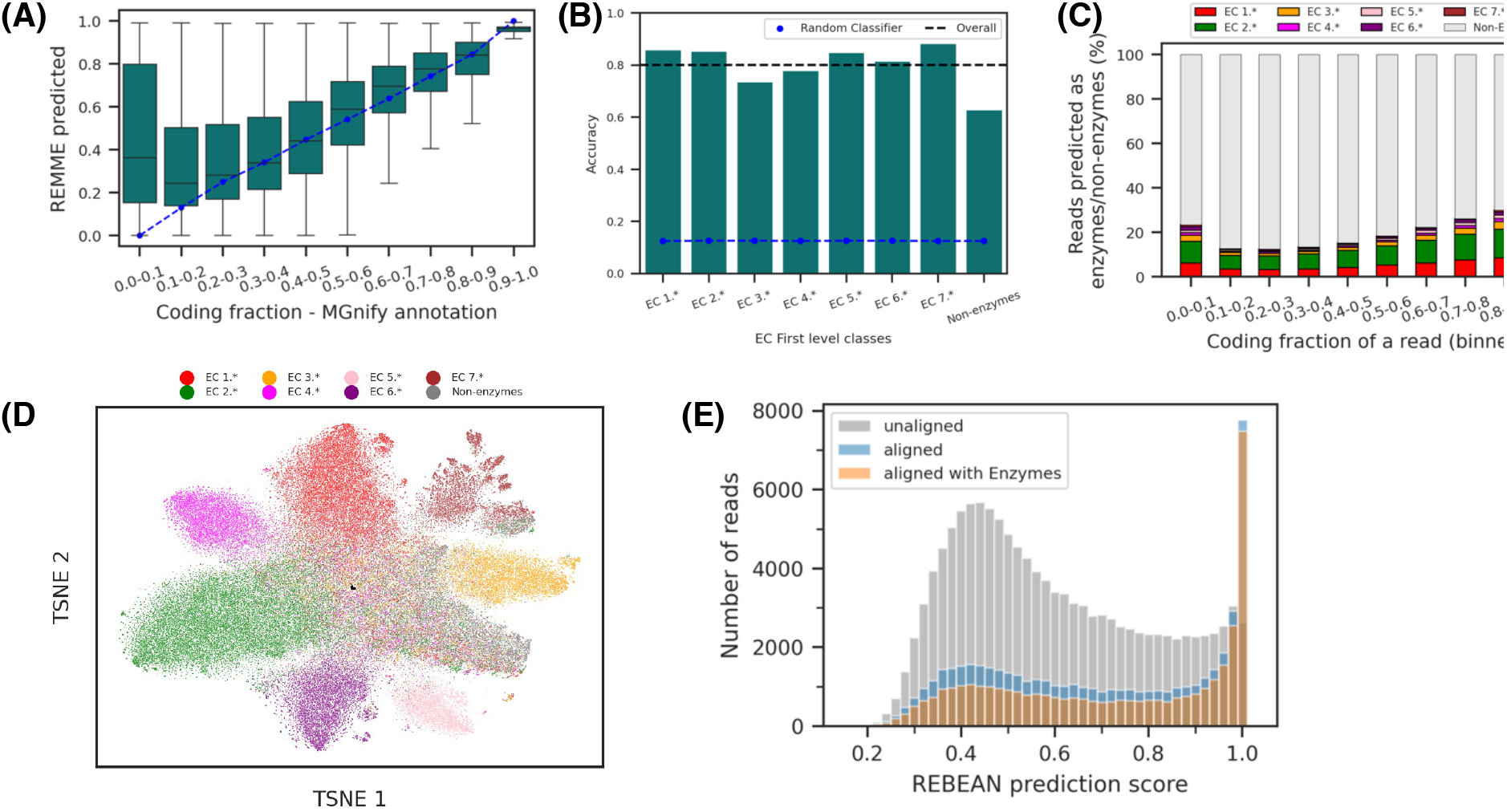
Training metagenomic read language models. **(A)** Across reads in REMME’s test set, the distribution of predicted, protein-coding nucleotide fractions (Y-axis) correlates with ground truth annotations (X-axis). **(B)** REBEAN’s test set prediction accuracy (**Eqn. 1**) highlights non-random performance across all enzyme classes (bars higher than the blue line), although some classes are deemed easier to label than others. **(C)** Distribution of reads in REMME’s test set across different fractions of protein-coding residues (X-Axis) vs. the number of these reads assigned to REBEAN-predicted EC classes (colors). Note that there is a trend from left to right, relating the coding fraction to the number of reads identified as coming from enzymes. For example, there are many more enzyme-like reads (∼47.4%) among those with a higher fraction of coding residues (0.9-1) than among those within the 0.1-0.2 coding fraction range (12.9%). **(D)** t-SNE projection of REBEAN embeddings of REMME’s test set, annotated with mi-faser-predicted EC classes (colors), illustrates REBEAN’s ability to differentiate read classes, **(E)** Distribution of REBEAN prediction scores for mi-faser predicted non-enzymatic reads in the test dataset that were REBEAN-predicted as enzymatic. Reads that do not align to any proteins in SwissProt (gray) tend to score lower than those that aligned with any protein (blue) or and aligned with enzymes (orange) at >30% sequence identity. A clear preference for higher scores of enzyme-aligned reads is visualized.

REMME predicted reading frames with an average accuracy of 48.6% and AUC of 0.789, i.e. not well but much better than a random four-class classifier of 25% accuracy. Note that annotation of CDSs and reading frames used in this study was produced by computational pipelines (Richardson et al., 2023) and, as such, could be incomplete or erroneous. Also note that the reading frame is not well defined for reads, i.e. sequence fragments, possibly shared by multiple CDSs with different reading frames on either forward or reverse strands. Despite these ambiguities, REMME distinguished reads which are part of CDSs (classes 1, 2 & 3) from non-transcribed reads with an accuracy of 88.5% (recall=94.2%, precision=92.7%; **Eqn. 4-5**).

In summary, REMME was pretrained, by enforcing relevant objectives, to understand the biological context of reads and did well in addressing the tasks it was assigned.

### Can we annotate enzyme-coding fragments in metagenomic data?

To answer this question, we fine-tuned REMME to build REBEAN – a model that predicts the Enzyme Commission (EC)(Tipton, 1994) first level classes (1: Oxidoreductases, 2: Transferases, 3: Hydrolases, 4: Lyases, 5: Isomerases, 6: Ligases & 7: Translocases) or a non-enzyme class of the proteins encoded by genes that gave rise to each read.

For the purposes of model training, read function annotations were generated by mi-faser (**Methods**), a high precision alignment-based read annotation method (Zhu et al., 2018). Relying on mi-faser annotations for training exposed REBEAN to a broader, real metagenome sequence space, instead of what can be gleaned from simulating reads from experimentally annotated reference databases. This decision also allowed us to train on reads from all genome regions, i.e. coding and non-coding, with no explicit biases.

Note that, somewhat counterintuitively, the task of labelling EC numbers at the first level is more challenging than predicting deeper EC levels, as deeper levels provide a more specific characterization of protein catalytic activity and typically correspond to only a handful of enzymes. For example, it is likely harder to find a unifying, broad, but unique, sequence or structure factor representing the first EC level, e.g. oxidoreductase activity (EC 1.*), than a specific factor explicitly capturing oxalate oxidase activity (EC number 1.2.3.4). Nevertheless, REBEAN attained consistently high accuracy (**Eqn. 1**) in classifying non-redundant reads (<80% sequence identity (Hauser et al., 2016)) across training, validation and test sets (81.7%, 81.0% and 80.6%, respectively; **Table S1** & **Figure 1C**). Furthermore, in testing, REBEAN attained an average multiclass Area Under the Curve (AUC, **Eqn. 9**) of 0.969 and recall of 80.1% with a precision of 83.6% (**Eqn. 4 & 5**). REBEAN’s test performance is far better than that of any random classifier (12.5% accuracy). Note that this performance reflects REBEAN’s ability to report mi-faser, not experimental, EC annotations.

To test whether pretraining helped training for EC prediction, we used REBEAN’s training protocol without REMME weights, instead initiating its base model with random assignments. This model reached a maximum accuracy of only 30% at the end of 200 epochs, illustrating the necessity of pre-training.

Despite the overall strong performance, REBEAN’s ability to identify non-enzymatic reads was relatively low (recall=50.6%). That is, REBEAN labelled as enzymatic half (176,254) of the test set reads, which mi-faser had not identified as such; **Figure 1D** illustrates that a portion of non-enzymatic reads’ REBEAN embeddings exists in the same space as enzymatic ones. We note again that mi-faser is a very precise (90% precision at fourth EC level) method of low recall (50%). We thus questioned whether these 176K reads could possibly indeed be enzymatic.

We further found that a third (50,756) of these reads aligned to SwissProt (Hauser et al., 2016; UniProt, 2023) proteins with >30% sequence identity. Three quarters (38,241) of these mapped to enzymes, highlighting alignment-based mi-faser method’s low recall and re-affirming reference-free REBEAN’s putatively high accuracy levels. Note, that reads aligned to SwissProt often attained higher REBEAN scores than those that did not align (**Figure 1E**). The remaining (126K) reads, predicted as enzymes by REBEAN, could represent yet-unseen enzymes or simply capture portions of gene sequences re-used in evolution across enzymatic and non-enzymatic sequences alike.

### REBEAN performs well in labelling functions on the basis of read sequences

To evaluate REBEAN’s performance against experimental, i.e. not mi-faser predicted, annotations, we generated a set of 525,775 fragments (length 200bp) from 4,295 enzyme-coding genes with experimental evidence of enzymatic activity (dataset **SPEnzset1, Table 1; Methods**). Each fragment, i.e. synthetic read, was labelled with the first level enzyme class of the parent gene. We evaluated REBEAN’s performance in annotating these reads’ enzyme classes (**Figure 2A, B**). At a threshold of 0.5, REBEAN attained an average recall=33.4%, precision=71.5%, and accuracy=88.1%; it attained an average Area Under the Receiver Operating Characteristics curve (AUROC, **Eqn. 9**) of 0.79, ranging from 0.73-0.90 across the seven enzyme classes (**Figure 2B** and **Table S1**).

**Figure 2:**
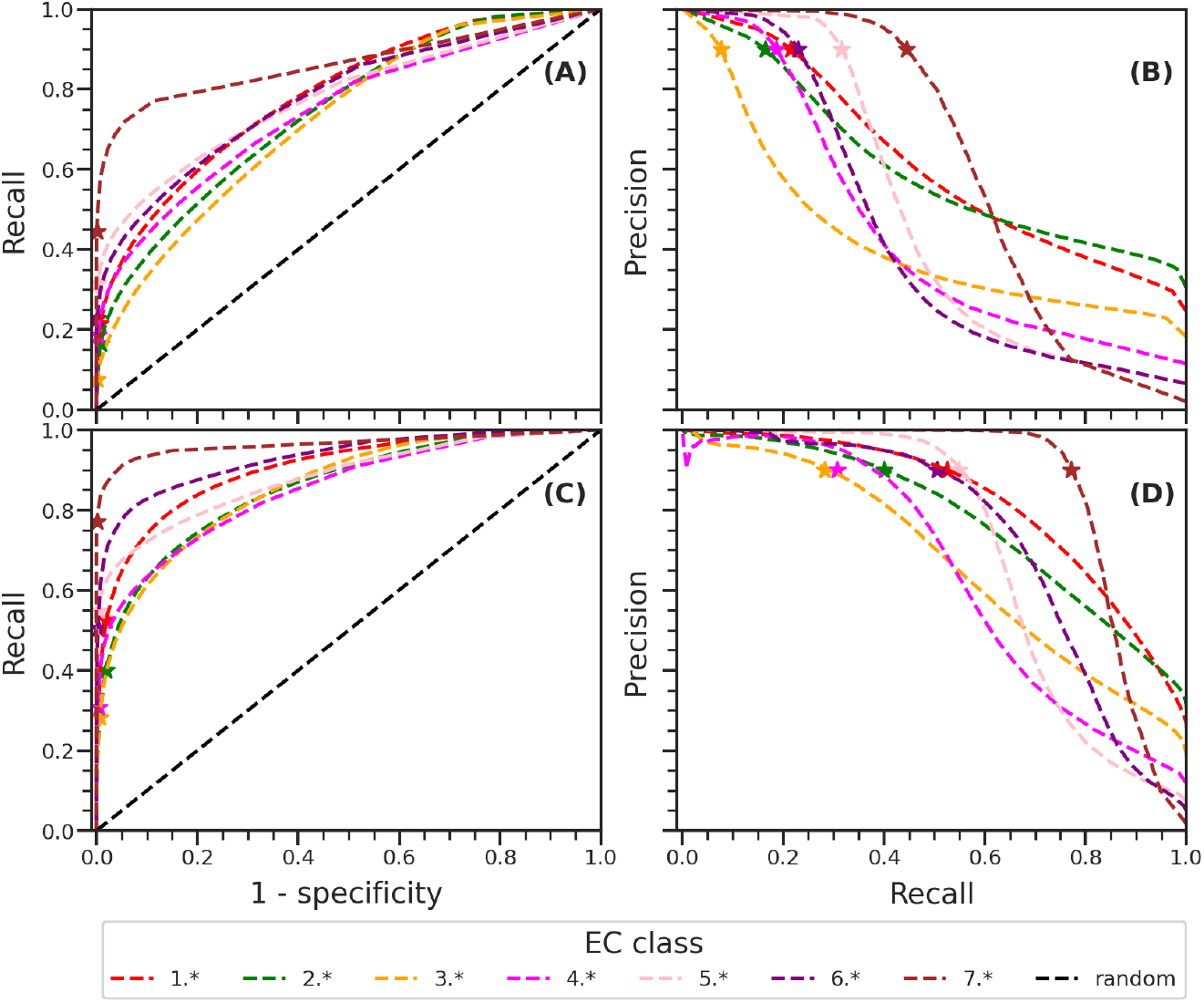
REBEAN synthetic read annotation. We extracted read-sized gene fragments from 4,295 prokaryotic enzymes and labelled them using REBEAN. **(A)** ROC and **(B)** PR curves demonstrate the performance of REBEAN in annotating reads with level 1 EC classes. **(C)** ROC and **(D)** PR curves represent performance of REBEAN in annotating complete proteins as an as average score of multiple reads with level 1 EC classes. The performance at chosen threshold corresponding to 90% precision is marked with ‘*’ on the curve.

Note that at 90% precision, REBEAN recovered, on average, only 23% (7.6% to 44.5%) of the reads from each enzyme class correctly; i.e., REBEAN attributed high prediction confidence to only a fraction of reads. Since each of the reads in this set is derived from an enzyme, REBEAN’s inability to assign high confidence to most reads suggested that it only learned to associate a specific segment in each gene with its molecular activity.

Three possible, non-exclusive, protein biology-driven explanations for this observation come to mind. (1) One is based on the fact that specific functional activities, e.g. catalysis and ligand binding, tend to occupy only *a small portion of the protein*, with the rest of the sequence providing structural support. In homologous enzymes, functional specificity is often tuned within a small protein segment, sometimes as short as a single residue (Allen & Whitman, 2021; Babbitt et al., 1996; Baier et al., 2016). Similarity between independently evolved, functionally similar enzymes is also often limited to a subset of residues, i.e. active-site convergence (Davidi et al., 2018). These functionally active protein sections are likely ubiquitous across enzyme classes (Bromberg et al., 2022). (2) Another possibility is that some *proteins may be multifunctional*, i.e. evolved to carry out different functions across different segments (Babbitt et al., 1996; Singh & Bhalla, 2020). (3) In the same vein, a read could be *part of multiple genes* encoding different regions of multiple proteins. The prokaryotic genome is densely packed, with an average of 88% of the genome encoding proteins (Chaumeil et al., 2019). In our analysis, as much as 7% of randomly generated reads could be expected to be shared by multiple genes (**Methods**).

### Improved Enzyme Annotation via Gene-Level Score Aggregation

We further tested REBEAN’s ability to label enzyme classes of complete genes on the basis of all corresponding read predictions. That is, we assigned each protein in our **SPEnzset1** (**Table 1**) the average score across the corresponding reads. Compared to read-based annotations, our model attained a much higher average AUROC of 0.89 (range: 0.86-0.97) and was able to recover 48% of the enzymes at 90% precision (**Figure 2C & D**).

We observed a strong positive correlation between the number of enzymes correctly classified and the number of reads sampled per gene. For example, the number of genes labelled correctly at 90% precision went up from 30% to 65% when the number of sampled reads increased from one to 50 reads per 1,000 basepairs of a gene. This finding strongly suggests that REBEAN would perform better for samples sequenced at higher depths (**Figure S2**).

We also assessed REBEAN’s ability to predict non-enzymes in our **SPset** (**Table 1**) – a problem at read level. Here, a whole protein could only be predicted to be non-enzymatic if all its reads were predicted non-enzymatic. REBEAN attained an AUROC of 0.83, in differentiating enzyme reads vs. putative non-enzyme reads, which is somewhat lower than its performance for labelling protein enzyme classes (∼0.89; **Table S1**). There are two potential explanations for this result: (1) some of the putative non-enzymes, as per their lack of annotation in SwissProt, may eventually be labelled as enzymes and (2) unlike enzyme classes, non-enzymatic proteins do not belong to a single class.

### REBEAN captures sequence fragments of functional importance

We investigated the biological relevance of REBEAN read prediction scores by exploring the distribution of catalytic and binding residues covered by these reads. We extracted the annotated functional site (catalytic and binding) residues from SwissProt proteins covered by our synthetic reads in **SPEnzset2** (UniProt, 2023). We observed an enrichment, i.e. a more frequent presence, of functional residues in reads with higher REBEAN scores. Reads predicted with a high prediction score, i.e. achieving 90% per-class precision (**Figure 2**), were 50% more likely to contain catalytic site residues than others (**Figure 3A**).

**Figure 3:**
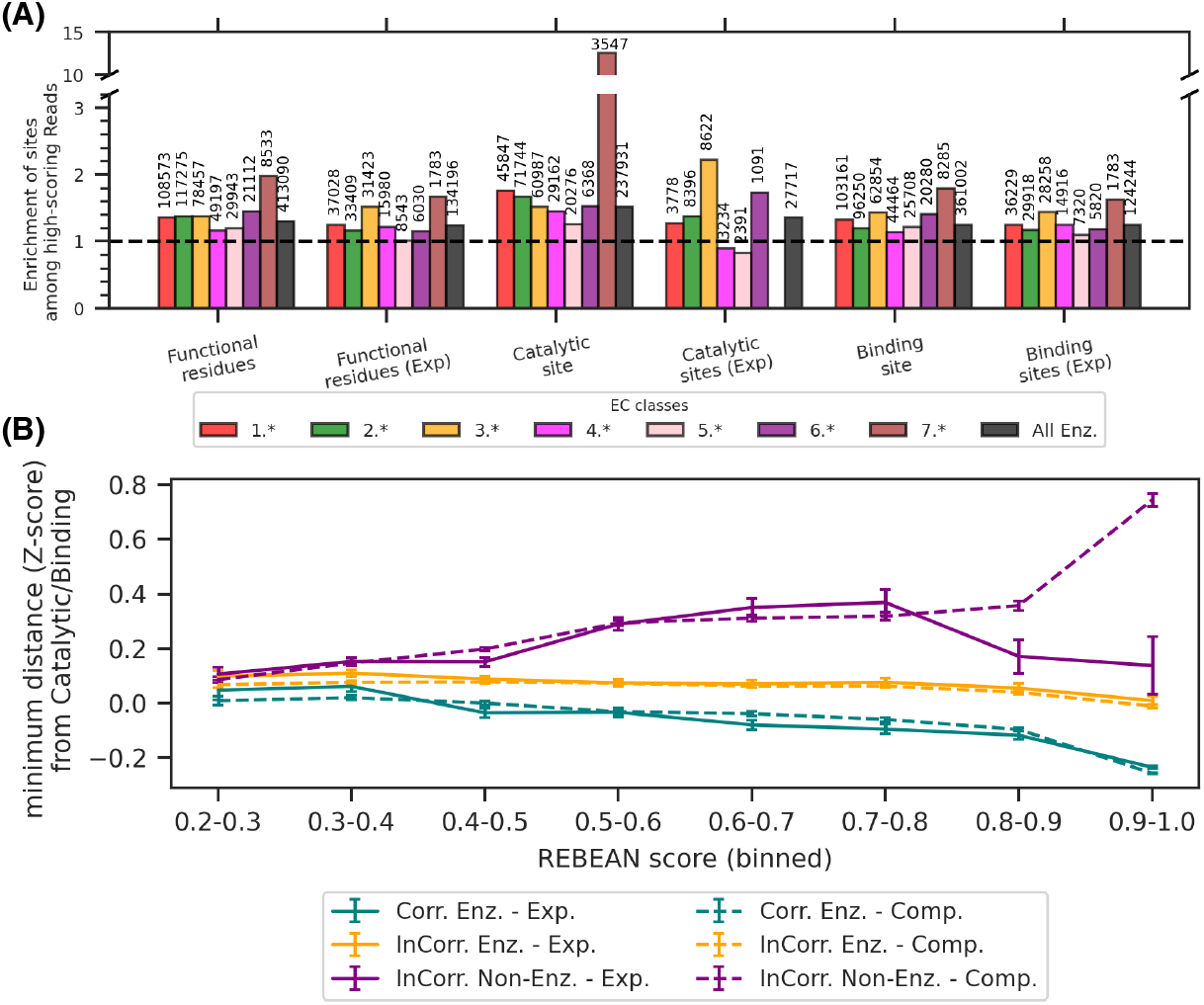
REBEAN high scoring enzymatic reads capture specifics of function. **(A)** Enrichment in residues annotated as active/catalytic and binding among reads predicted with high confidence across enzyme classes (colors). All bars are higher than 1 (dashed line), indicating enrichment vs. numbers (bar height) expected for all reads. Numbers on top of the bars indicate the number of reads for which the enrichment was calculated. **(B)** Normalized spatial (3D) distance of the reads from functional residues (active and binding) computed using experimental (solid lines) and AlphaFold2 predicted protein structures (dashed lines). Reads correctly predicted by REBEAN to their enzyme classes (Corr. Enz., teal) are structurally closer (lower Y-axis) to functional residues vs. reads incorrectly labelled as non-enzymatic (InCorr. Non-Enz., purple) or assigned a different enzyme class (orange, InCorr.Enz.)

As function site annotations in databases are often incomplete, we further investigated the spatial distribution of read-encoded amino acid residues in the vicinity of the functional sites in protein structures. We calculated the minimum distance from the read-encoded amino acids to the structurally closest functional site residues. The calculated distance for each read was normalized across all reads from a given gene to account for variable gene lengths. Again, we observed that translations of high scoring reads were significantly closer in 3D space to functional residues (**Figure 3B**).

### REBEAN identifies orthologs without relying on sequence similarity

We extended our analysis of REBEAN embeddings to compare reads sampled from OrthoDB (Kuznetsov et al., 2023) genes, representing orthologous and non-orthologous gene pairs across different taxonomy levels, i.e. genus, family, order, class and phylum (**Orthoset1, Table1; Methods**). For each gene pair, we calculated the average cosine similarity between read embeddings from each of the two genes (**Eq. 10**). As expected from our earlier work (Hoarfrost et al., 2022), we observed a significant distinction in embedding similarity between reads of orthologous vs. non-orthologous gene pairs at all five taxonomic levels (p-value < 1E-32; **Figure 4**). We also noted a small but significant drop in embedding similarity between reads from orthologous gene pairs in successive taxonomic levels from genus to phylum (*p-value* < 4E-4 - 1E-102).

**Figure 4:**
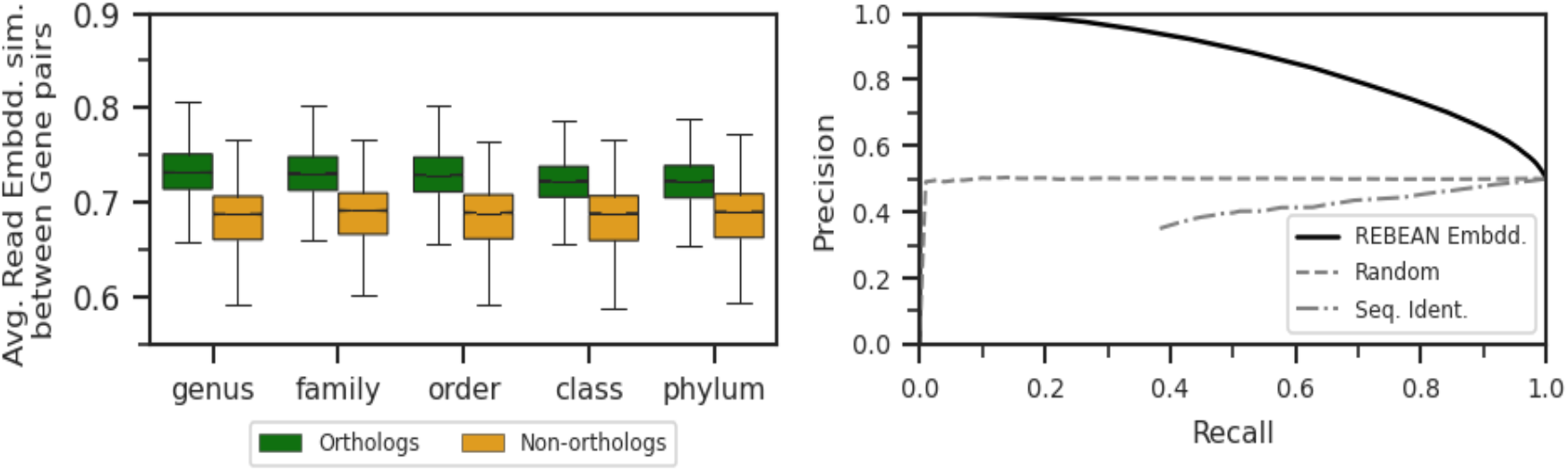
Fragments from Ortholog gene pairs share embedding space. **(A)** Distribution of average embedding similarity between reads from orthologous and non-orthologous gene pairs. **(B)** PR curve for the prediction of orthologous vs. non-orthologous gene pairs using embedding similarity.

We further asked the reverse question: given a particular average read embedding similarity, is the gene pair more likely to be from an orthologous or a non-orthologous set? Embedding similarities were accurate (74.6%) in making this identification (AUROC= 0.84, AUPRC=0.86; **Table 2**) – a performance significantly better than random (50%) or that of MMseqs-derived (Hauser et al., 2016) sequence identities of gene pairs (50*%*). As expected, we again observed better performance for lower taxonomic levels; e.g. for gene orthologous pairs at genus level, the accuracy was 79.6% (AUROC=0.88, AUPRC=0.90), while at phylum level the accuracy dropped to 70% (AUROC=0.81, AUPRC=0.82). Note that REMME (instead of REBEAN) embeddings attained a similar performance at this task.

**Table 2:** Predicting orthologs gene pairs across taxa.

| Taxa | Method | AUC ROC | AUC PR | F1 max | threshold<sup>1</sup> | Recall | Precision | Accuracy |
| --- | --- | --- | --- | --- | --- | --- | --- | --- |
| Genus | REBEAN | 0.880 | 0.895 | 0.799 | 0.707 | 0.830 | 0.770 | 0.791 |
| Family | REBEAN | 0.851 | 0.866 | 0.772 | 0.703 | 0.846 | 0.709 | 0.750 |
| Order | REBEAN | 0.854 | 0.869 | 0.773 | 0.706 | 0.810 | 0.739 | 0.762 |
| Class | REBEAN | 0.824 | 0.835 | 0.753 | 0.699 | 0.820 | 0.697 | 0.732 |
| Phylum | REBEAN | 0.808 | 0.82 | 0.742 | 0.693 | 0.861 | 0.651 | 0.700 |
| All | REBEAN | 0.843 | 0.858 | 0.766 | 0.701 | 0.837 | 0.705 | 0.744 |
| All | Seq. Similarity | 0.319 | 0.526 | 0.667 | 0 | 1 | 0.5 | 0.5 |

<sup>1</sup>The threshold corresponds to the maximum F1 score.

Instead of focusing on whole genes/proteins, we aligned a set of reads (**Orthoset2**) to find the correlation between read sequence identity (**Methods**) and REMME embedding similarity (**Eqn. 11**). Correlation was fairly low for read pairs of (1) the same gene (Pearson r=0.08), (2) different genes (0.35), (3) genes of the same orthologous group (0.09-0.16 across taxa), or (4) genes of different orthologous groups (0.14-0.30). REBEAN read embedding similarities were also indifferent to sequence identity across gene pairs (|r| < 0.07). In other words, read embeddings carried different information than sequence identity.

We note, however, that fine-tuning our models to predict enzymatic function reduced the correlation between embedding similarity and sequence identity. That is, REMME (pretrained LM) read embedding similarities were somewhat more like sequence identities than REBEAN (fine-tuned for EC prediction) read embedding similarities. REMME is a DNA language model trained to encode information embedded in sequence and hence is expected to capture sequence identity. It is thus useful in sequence encoding for, among many applications, identifying coding regions and estimating the likelihood of a given DNA read coming from a specific genome (**Figure S3**). However, tuning the model to predict function loosens the need for explicit sequence representation.

### REBEAN can be used to discover novel enzymes from metagenomes

REBEAN’s ability to capture function without requiring sequence similarity can help in identifying proteins that carry out known functions in novel ways. Here, we aimed to evaluate REBEAN’s ability to mine potential novel oxidoreductases from a metagenomic set of reads. Oxidoreductases are broad class of enzymes that catalyse redox reactions by facilitating transfer of electrons. Oxidoreductases are ubiquitous as they are part of every biological energy production mechanism (Hay Mele et al., 2023; Kim et al., 2013) and are likely to have appeared on the scene early in history of life on Earth (Bromberg et al., 2022).

For this analysis, we compiled a synthetic read dataset randomly sampled from the assembled sequences of metagenome-assembled genomes (MAGs) in MGnify (Richardson et al., 2023) (Gurbich et al., 2023) (**MarineMGset, Methods**). REBEAN predicted that approximately 10% (12,861,390) of these reads as enzymatic with a high confidence score of >0.9; of these, a quarter (3,019,892 reads) were deemed oxidoreductases (**Figure S4**). Note that almost all (98.7%) of the predicted oxidoreductase reads mapped to MGnify-labelled genes (1,126,995 genes) (Richardson et al., 2023), i.e. this prediction rarely tagged non-coding regions.

In search for truly novel enzymes, we retained only the reads that mapped to genes without any existing MGnify annotations (COG, EC, KEGG, Pfam); due to MGnify filters, for many of the MAG genes the corresponding protein sequences and annotations, were not available. This analysis retained a third of the genes (407,428 genes; 670,637 reads) distributed across all 3,820 MAGs. We further analyzed the available protein sequences encoded by these genes (275,290 of 407,428).

We extracted Pfam domains of these sequences using HMMER (Eddy, 2011). Curiously, although MGnify was missing this annotation for 234,025 proteins, our analysis revealed their 6,883 unique Pfams (E-value < 1E-3, excluding 1,926 DUFs), collectively occurring 740,053 times. To evaluate the significance of Pfam assignments in predicting enzymatic activity — particularly oxidoreductase function — we compared the Pfam distribution within our dataset to that of oxidoreductases in SwissProt (**Table S3**). We observed a significant enrichment of Pfams associated with SwissProt-annotated oxidoreductases in our dataset (Odds ratio = 2.06, Hypergeometric p-value < 1E-32; **Methods**).

We clustered (Hauser et al., 2016) the 275K protein sequences at 30% sequence identity, yielding 51,326 representative sequences. Aligning these against SwissProt (≤30% sequence identity), we identified 39,617 putatively novel proteins. For 32,030 of these proteins (length 100 to 800 residues), we generated structure predictions using ESMFold (Lin et al., 2023). Pfam oxidoreductase analysis, as above, confirmed our predictions (Odds ratio = 2.43, **Table S3**). Note that these proteins were classified based on the prediction of a single read with oxidoreductase activity. Integrating predictions from all reads corresponding to a protein previously demonstrated higher accuracy (**Figure 2C & D**). Using this refined approach here, we identified 4,901 proteins predicted to function as oxidoreductases at a 90% precision threshold. These proteins exhibited even greater enrichment in oxidoreductase-associated Pfams (Odds ratio = 3.66). These results suggest that REBEAN can, no reference needed, reveal novel proteins that carry out known functions.

To examine the structural similarity of the read-based group of REBEAN-identified proteins (32K) with existing oxidoreductases, we aligned their structures against the PDB database using FoldSeek (van Kempen et al., 2024) to find that 464 (1.4% of the structures) matched 1,698 PDB entries (TM ≥ 0.9). Roughly half (938) of these PDBs lacked an EC annotation. Among the remaining 760 enzymes, half were classified as oxidoreductases (EC 1; 373 structures, 49% of enzymes), confirming REBEAN’s ability to label enzyme classes.

The remaining enzyme matches comprised 143 transferases (19%; EC 2), 128 hydrolases (17%; EC 3), 28 lyases (4%; EC4), 86 isomerases (11%; EC 5) and 2 ligases (EC 6). While these matches suggest REBEAN’s error in function labelling, additional considerations may be at play. For example, the top two non-oxidoreductase EC classes matched by this alignment were 2.7.7.7 (DNA polymerase, 42 structures) and 5.3.1.5 (Xylose Isomerase, 31 structures). DNA polymerase serves a critical but a non-redox function. However, these enzymes, like oxidoreductases, bind cations and nucleotides – a functionally crucial activity (Hay Mele et al., 2023; Selles Vidal et al., 2018; Steitz, 1999). EC class 5.3.1.5 is also like EC 1’s; it is an isomerase subclass of intramolecular oxidoreductases (EC 5.3) that carries out oxidation and reduction within a molecule. We thus suggest that, at least in this case 19% (72 of 387) of the REBEAN errors may be driven by functional similarity of enzyme regions due to evolutionary and/or molecular activity constraints. For other putative false positive labels while some overlap with other enzyme classes may indeed be an error, it is also possibly a consequence of the evolutionary promiscuity and functional crossover observed among enzymes (Martinez Cuesta et al., 2015).

### Interpreting REBEAN predictions through Embeddings

What did the model learn? To answer this question, we computed read-pair Euclidean similarity (**Eqn. 10**) and cosine similarity (**Eqn. 11**) between confidently predicted reads with REBEAN score above 0.90. For both metrics, we then computed the average similarity (**Eqn. 12-13**) of each read with other reads in our dataset of 54,115 reads generated from 4,295 enzyme genes (**SPEnzset2, Table 1**). Note that both cosine and Euclidean similarity metrics maintained similar trends, so from here on we will only describe the Euclidean similarity results.

The average similarity of a read with other reads sampled from the same gene was 0.64±0.06 and varied widely from 0.37 to 1.0 (**Figure 5A**) indicating contextual differences of various gene regions. On the other hand, similarities of read embeddings from different genes were only slightly lower (0.59±0.03), irrespective of their EC class (range: 0.30 to 0.64). These observations reinforce the conclusion that much of any given gene sequence is likely shared between enzymes for e.g. structural or evolutionary reasons, with only a small fraction being specific to a particular function.

**Figure 5:**
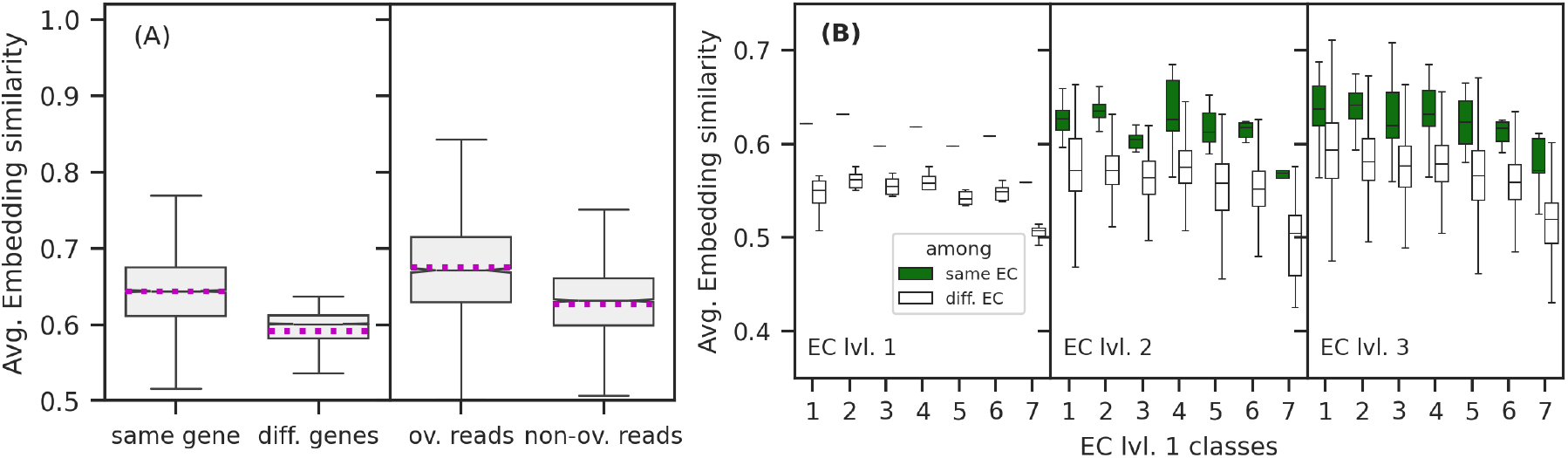
REBEAN embedding similarity captures sequence and functional similarity. **(A)** Distribution of average embedding similarities of each read vs. reads from the same gene or different genes, overlapping or not, for reads predicted with score above 0.90. The mean embedding similarity for each distribution is marked by the magenta line **(B)** Distribution of average embedding similarities of each read across the seven EC classes with reads from other enzyme classes at different EC levels (first, second and third).

Similarity between overlapping reads within a gene was higher than that of non-overlapping reads (**Figure 5A**), i.e. as expected, reads that share a portion of the same exact sequence within a gene tend to have higher embedding similarity. Note that, as we had demonstrated with the ortholog dataset, we did not observe any correlation between embedding similarity and sequence identity of read pairs from different genes (Hauser et al., 2016). Together, these observations suggest that the model indeed learned functional, rather than solely sequence-based, signatures encoded in genomic data. We thus probed deeper into the question: how much function did the model learn?

We computed the embedding similarity between all pairs of reads within same EC class at first, second, and third EC levels (e.g. EC 1.*, 1.1.*, and 1.1.9.*; **Figure 5B**). At all three levels, reads from same enzyme classes were more similar than reads from different classes, although the difference was more pronounced for higher EC levels (first>second>third level). Note, however, that the embedding similarities were higher within lower EC levels (first<second<third). The average similarity varied across enzyme classes (EC1-7), with translocase (EC 7.*) reads tended being least similar. This observation is in line with the fact that this class comprises subgroups that were historically part of EC designations of oxidoreductases, hydrolases, and lyases (McDonald & Tipton, 2023).

### Comparing REBEAN predictions with other metagenome annotation methods

We compared REBEAN predictions for 8 million reads from 18 metagenomic samples (**Table 1: ExtremeMGset**) from three aquatic environments (hydrothermal vents, hypersaline and salt crystallizer ponds) against annotations of other methods, i.e. mi-faser, HUMAnN3, Carnelian and LookingGlass (**Methods**) (Beghini et al., 2021; Hoarfrost et al., 2022; Nazeen et al., 2020; Zhu et al., 2018). The selection of extreme environments for this evaluation is in line with the desire for novel function discovery. Furthermore, these environments provide a microbial diversity profile distinct from those typically represented in reference databases, which are typically used by alignment-based annotation tools.

Due to the lack of ground-truth labels, we assessed each method at its default cutoff based on its ability to annotate reads (coverage) and consistency of annotation with other tools, i.e. mutual agreement in annotated enzymatic class. As a baseline, we aligned the 8M reads to UniRef90 (release 201901b) using DIAMOND and labelled any read that matched with an enzyme as enzymatic read of the enzyme class (**Methods**, (Buchfink et al., 2021)). Note that default cutoffs for all methods balance precision of annotation against coverage, so a larger number of reads could be annotated by most at lower accuracy. However, method optimization and, thus, this discussion are beyond the scope of this manuscript.

Across all environments REBEAN, at its default 90% precision cutoff, consistently identified 3-to 6-fold more enzymatic reads than alignment-based tools. The alignment-based tools were, at best, only able to label 2% of the reads as coming from enzyme-coding genes (**Figure 6A & B, Figure S5**). For example, HUMAnN, the state-of-the-art pipeline incorporating DIAMOND, annotated a mere 0.2% of reads from extreme environments. This highlights a significant limitation of methods that rely on sequence similarity to reference databases.

**Figure 6:**
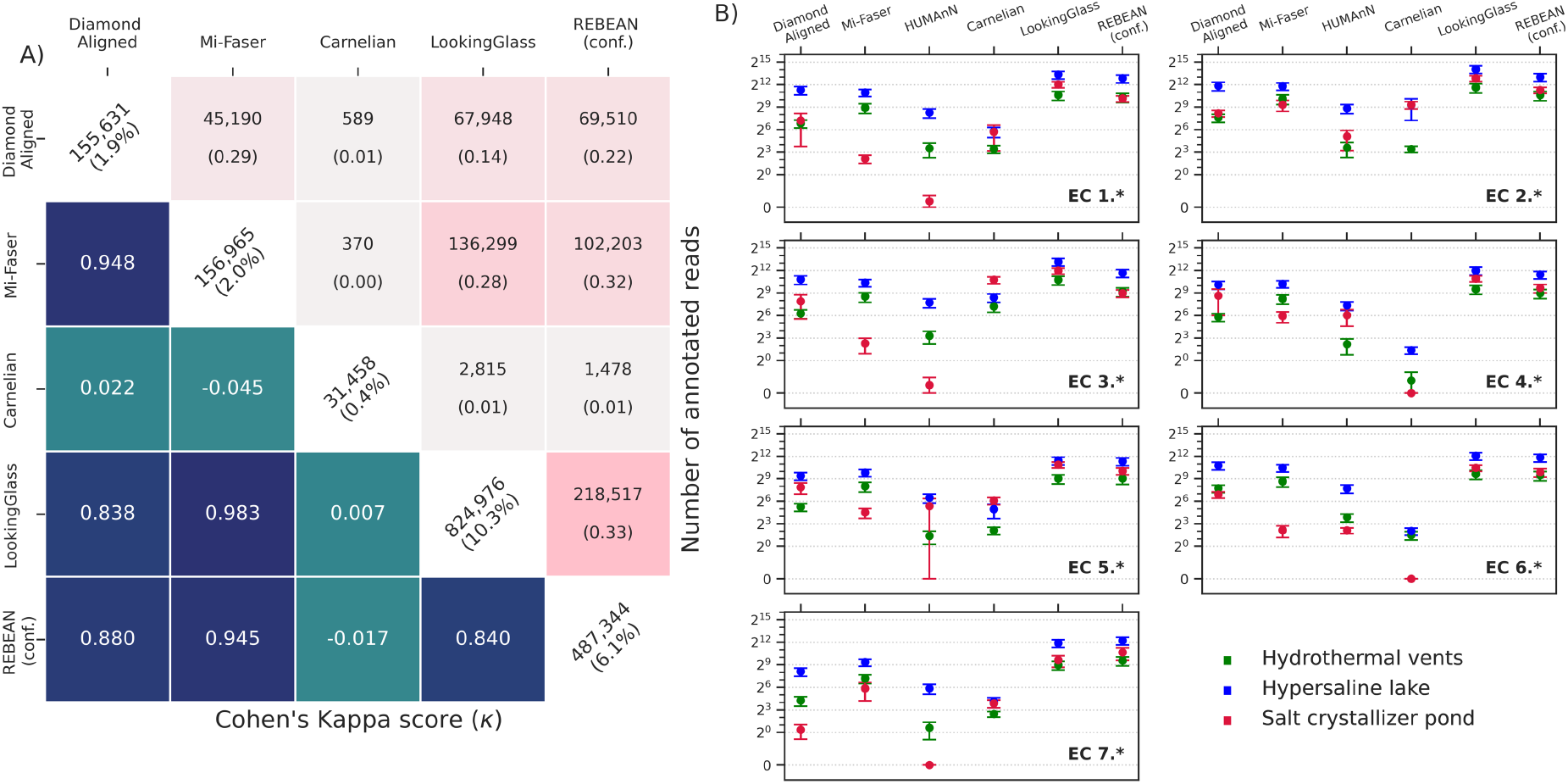
Functional annotation of extremophilic microbiomes. **(A)** Cohen’s kappa score (below the diagonal) quantifies the agreement of predicted first level EC class labels between different tools, summed across all samples. Higher kappa scores indicate stronger agreement than expected by chance. The counts and fractions of jointly annotated reads between each pair of tools are shown above the diagonal. Diagonal values indicate the number of annotated reads out of the 8M reads annotated by each tool. **(B)** Read counts annotated to the seven first level EC classes across the 25 samples from three environments: hydrothermal vents (green), hypersaline (blue) and salt crystallizer ponds (crimson). The y-axis is log_2_-transformed to visualize differences in read counts across samples. REBEAN and LookingGlass consistently annotate at least three-fold more reads than other tools.

In contrast, reference-free machine learning tools can overcome this restriction if they learn to generalize functional sequence signatures beyond sequence similarity; that is, they can, in theory, annotate a much higher proportion of reads. Indeed, REBEAN (at a high 90% precision threshold) was able to annotate 6.1% of the reads, (36.4% without any restriction), while LookingGlass identified 10.3% of reads as enzymatic. Note that machine-learning-based Carnelian annotated only 0.4% of reads, a number that may indicate method dependence on threshold choice.

Devoid of a ground truth for direct method performance assessment, we computed Cohen’s Kappa scores (ranges from -1 to 1) comparing all tools to measure consistency of the annotations (**Figure 6A**). Note that HUMAnN was excluded from this analysis as its gene-family annotations are not directly comparable to read-level annotations. REBEAN’s annotations aligned well with both mi-faser and DIAMOND (Cohen’s kappa =0.945 and 0.880, respectively), indicating likely correctness of jointly labelled reads. Even without the class-specific labels, REBEAN shares the highest jointly annotated enzymatic reads with other tools: diamond-alignment (22%), mi-faser (32%), and LookingGlass (33%).

Taken together, these findings illustrate REBEAN’s ability to expand accurate description of metagenome functionality, i.e. increase the number of annotated reads.

### Addressing the need to step up tool development

Accurate functional annotation of metagenomes is an important step in understanding microbial communities. However, discovery and identification of novel functions in metagenomic data requires innovative approaches that transcend traditional homology-based methods (Prabakaran & Bromberg, 2025). For metagenomic tools to effectively aid in discovery, they must be both robust and highly accurate.

Recent advances in language model development have accelerated the field of protein structure and function prediction (J. Jumper et al., 2021; Lin et al., 2023) and we believe that a similar leap in metagenomic functional annotation is essential.

We believe that tools like REMME and REBEAN can pave the way for this transformation. By capturing biologically relevant insights, REBEAN has shown considerable potential for the discovery of truly novel, i.e. sequence and structure dissimilar, means of carrying out enzymatic activity. This capability is crucial as the field moves toward exploring the uncharted territory of novel enzymes.

### Summary of Findings

The increasing accumulation of metagenomic samples presents both new challenges and opportunities for tool development. To address this, we have developed REMME, a robust foundational DNA-language model tailored to capture and interpret biological context encoded within metagenomic reads. REMME’s versatility makes it valuable for diverse research applications including generating read embeddings for machine learning applications, clustering reads for comprehensive ecological and evolutionary studies and, also, building fine-tuned models like REBEAN for specific downstream tasks.

REBEAN, a method for predicting enzymatic functionality from read information alone, was trained using millions of diverse metagenomic reads. Unlike alignment-based methods, REBEAN can discover novel enzymatic sequences. Moreover, though never explicitly trained to do so, its predictions highlight functionally significant residues within a given read. Our extensive analysis highlights REMME’s and REBEAN’s potential for metagenomic read annotation and the discovery of novel enzymes – a route in metagenomic exploration that has not yet been attempted by other computational techniques.

## Supporting information

SOM

## Acknowledgments

This work was supported by the NASA Astrobiology Institute Grant Number: 80NSSC18M0093. It was also supported by the NSF (National Science Foundation) awards #2310114.

## Author contributions

Conceptualization: YB; Methodology: RP, YB; Investigation: RP; Visualization: RP; Supervision: YB; Writing and revisions: RP, YB;

## Data and materials availability

All data are available in the main text or the supplementary materials. The models and supplementary scripts are publicly available at https://bitbucket.org/bromberglab/rebeanpkg. In addition, REBEAN is accessible as a free web service at https://services.bromberglab.org/rebean/.

