## Supplementary material for "Deciphering enzymatic potential in metagenomic reads through DNA language models": SOM

**Prabakaran R[1,2*], Bromberg Y[1,2*]**

[1] Department of Biology, Emory University, Atlanta, GA 30322, USA
[2] Department of Computer Science, Emory University, Atlanta, GA 30322, USA

### Supplementary Information

#### Table S1: Performance of REBEAN in predicting enzyme class from reads

| CLASS | DATASET | PRECISION | RECALL | F1-SCORE | AUC | Count of reads |
|---|---|---|---|---|---|---|
| **1** | Training | 0.821 | 0.868 | 0.844 | 0.978 | 2,418,548 |
| **2** | Training | 0.836 | 0.861 | 0.848 | 0.971 | 3,538,693 |
| **3** | Training | 0.868 | 0.745 | 0.802 | 0.970 | 1,540,732 |
| **4** | Training | 0.924 | 0.790 | 0.852 | 0.980 | 1,198,884 |
| **5** | Training | 0.882 | 0.857 | 0.869 | 0.985 | 794,906 |
| **6** | Training | 0.933 | 0.825 | 0.876 | 0.983 | 1,431,171 |
| **7** | Training | 0.968 | 0.889 | 0.927 | 0.993 | 1,046,580 |
| **8** | Training | 0.514 | 0.641 | 0.571 | 0.910 | 1,709,930 |
| **AVERAGE** | Training | 0.843 | 0.809 | 0.823 | 0.971 | 13,679,444 |
| **WEIGHTED** | Training | 0.827 | 0.814 | 0.818 | 0.969 | 13,679,444 |
| **1** | Validation | 0.812 | 0.860 | 0.835 | 0.975 | 268,728 |
| **2** | Validation | 0.827 | 0.855 | 0.841 | 0.969 | 393,188 |
| **3** | Validation | 0.858 | 0.734 | 0.791 | 0.967 | 171,192 |
| **4** | Validation | 0.919 | 0.779 | 0.843 | 0.977 | 133,209 |
| **5** | Validation | 0.876 | 0.847 | 0.861 | 0.983 | 88,323 |
| **6** | Validation | 0.927 | 0.817 | 0.868 | 0.981 | 159,019 |
| **7** | Validation | 0.966 | 0.882 | 0.922 | 0.992 | 116,287 |
| **8** | Validation | 0.505 | 0.633 | 0.562 | 0.907 | 189,992 |
| **AVERAGE** | Validation | 0.836 | 0.801 | 0.815 | 0.969 | 1,519,938 |
| **WEIGHTED** | Validation | 0.820 | 0.805 | 0.810 | 0.967 | 1,519,938 |
| **1** | Test | 0.812 | 0.861 | 0.836 | 0.976 | 671,817 |
| **2** | Test | 0.826 | 0.854 | 0.840 | 0.969 | 982,974 |
| **3** | Test | 0.858 | 0.737 | 0.793 | 0.968 | 427,987 |
| **4** | Test | 0.917 | 0.781 | 0.843 | 0.977 | 333,023 |
| **5** | Test | 0.875 | 0.850 | 0.862 | 0.983 | 220,812 |
| **6** | Test | 0.926 | 0.817 | 0.868 | 0.981 | 397,546 |
| **7** | Test | 0.966 | 0.884 | 0.923 | 0.992 | 290,722 |
| **8** | Test | 0.506 | 0.629 | 0.561 | 0.906 | 474,983 |
| **AVERAGE** | Test | 0.836 | 0.801 | 0.816 | 0.969 | 3,799,864 |
| **WEIGHTED** | Test | 0.819 | 0.806 | 0.810 | 0.967 | 3,799,864 |

### Table S2: List of 360 experimental runs selected for REBEAN training

| | | | | | | |
|---|---|---|---|---|---|---|
| ERR058011 | ERR058011 | ERR058014 | ERR058017 | ERR058017 | ERR058019 | ERR058021 |
| ERR058022 | ERR058023 | ERR058023 | ERR058024 | ERR058024 | ERR058025 | ERR058026 |
| ERR058029 | ERR058029 | ERR058031 | ERR058031 | ERR058034 | ERR058035 | ERR058035 |
| ERR058036 | ERR058036 | ERR058037 | ERR058038 | ERR058039 | ERR058040 | ERR058042 |
| ERR058043 | ERR058045 | ERR058046 | ERR058046 | ERR058047 | ERR058047 | ERR058051 |
| ERR058053 | ERR058054 | ERR058055 | ERR058057 | ERR058058 | ERR205703 | ERR205705 |
| ERR205717 | ERR205719 | ERR205720 | ERR2102868 | ERR2102871 | ERR2102875 | ERR2102877 |
| ERR2102883 | ERR2102890 | ERR2102891 | ERR2102894 | ERR2102900 | ERR2102910 | ERR2102916 |
| ERR2102917 | ERR2102919 | ERR2102922 | ERR2102931 | ERR2102946 | ERR2102946 | ERR2102949 |
| ERR2102951 | ERR2102951 | ERR2102956 | ERR2102956 | ERR2102964 | ERR2102970 | ERR2102971 |
| ERR2102975 | ERR2102977 | ERR2102986 | ERR2102987 | ERR2102988 | ERR2102989 | ERR2102995 |
| ERR2102996 | ERR2102998 | ERR2103000 | ERR2103013 | ERR2103018 | ERR2103024 | ERR2103029 |
| ERR2103032 | ERR2172154 | ERR2172155 | ERR2172165 | ERR2172173 | ERR2172173 | ERR2172174 |
| ERR2172174 | ERR2172175 | ERR2172185 | ERR2172186 | ERR2172206 | ERR2172209 | ERR2172215 |
| ERR2172230 | ERR2172243 | ERR2172245 | ERR2172245 | ERR2172248 | ERR2172248 | ERR2172249 |
| ERR2172250 | ERR2172251 | ERR2215427 | ERR2215432 | ERR2215436 | ERR2215437 | ERR2215444 |
| ERR2215447 | ERR2215454 | ERR2215455 | ERR2222752 | ERR2222756 | ERR2222757 | ERR2222758 |
| ERR2528566 | ERR2528572 | ERR2528614 | ERR2529233 | ERR2599709 | ERR2599710 | ERR2599718 |
| ERR2599725 | ERR2599728 | ERR2599731 | ERR2599744 | ERR2601411 | ERR2601413 | ERR2601418 |
| ERR2601420 | ERR2601430 | ERR2601441 | ERR2601443 | ERR2601450 | ERR2601453 | ERR2601455 |
| ERR2601462 | ERR2601481 | ERR2601482 | ERR2601482 | ERR2601484 | ERR2601490 | ERR2601490 |
| ERR2601504 | ERR2601505 | ERR2601506 | ERR2601516 | ERR2601518 | ERR2601521 | ERR2601527 |
| ERR2601529 | ERR2601532 | ERR2617118 | ERR2617123 | ERR2617128 | ERR2617133 | ERR2617134 |
| ERR2617135 | ERR2617144 | ERR2617153 | ERR2617154 | ERR2617154 | ERR2617156 | ERR2617157 |
| ERR2617178 | ERR2696813 | ERR2709717 | ERR2709717 | ERR2709720 | ERR2709730 | ERR2709733 |
| ERR2709739 | ERR2709748 | ERR2709750 | ERR2709761 | ERR2709767 | ERR2709768 | ERR2709769 |
| ERR2709786 | ERR2709789 | ERR2709792 | ERR2709797 | ERR2709815 | ERR2777825 | ERR2777828 |
| ERR2777829 | ERR2777831 | ERR2777833 | ERR2777835 | ERR2777836 | ERR2777849 | ERR2777850 |
| ERR2777852 | ERR2777852 | ERR2777855 | ERR2808649 | ERR2808650 | ERR2808651 | ERR2808652 |
| ERR2808653 | ERR2808654 | ERR2808657 | ERR2808662 | ERR2808664 | ERR2808665 | ERR2808666 |
| ERR2808667 | ERR2814646 | ERR2814647 | ERR2814649 | ERR2814651 | ERR2814656 | ERR2814660 |
| ERR2814662 | ERR2814665 | ERR2814666 | ERR2814988 | ERR2814991 | ERR2814994 | ERR2814996 |
| ERR2814997 | ERR2814998 | ERR2815000 | ERR2815000 | ERR2815001 | ERR2815002 | ERR2815004 |
| ERR2815007 | ERR2815007 | ERR2816120 | ERR2816127 | ERR2816129 | ERR2816139 | ERR2816142 |
| ERR2816143 | ERR2816148 | ERR2816150 | ERR2816154 | ERR2816156 | ERR2816159 | ERR2816164 |
| ERR2816165 | ERR2816167 | ERR2816168 | ERR2816184 | ERR2816185 | ERR2816187 | ERR2816188 |
| ERR2816191 | ERR2816192 | ERR2816208 | ERR2816208 | ERR2816215 | ERR2816219 | ERR2816226 |
| ERR2816227 | ERR2985257 | ERR2985263 | ERR2985266 | ERR2985270 | ERR2985271 | ERR2985273 |
| ERR3256330 | ERR3256341 | ERR3256346 | ERR3256348 | ERR3256371 | ERR3256375 | ERR3256378 |
| ERR3256383 | ERR3256385 | ERR3335351 | ERR3361769 | ERR3361773 | ERR3361775 | ERR3361777 |
| ERR3361779 | ERR3361780 | ERR3361782 | ERR3361784 | ERR3361785 | ERR3361788 | ERR3361789 |
| ERR3361792 | ERR3361793 | ERR3361794 | ERR3361795 | ERR3361798 | ERR3361800 | ERR3361804 |
| ERR3361812 | ERR3361813 | ERR3361815 | ERR3367835 | ERR3367842 | ERR3367843 | ERR3367867 |
| ERR3367867 | ERR3367870 | ERR3367874 | ERR3367874 | ERR3367886 | ERR3367890 | ERR3367891 |
| ERR3378956 | ERR3378958 | ERR3378959 | ERR3378966 | ERR3378983 | ERR3378987 | ERR3379002 |
| ERR3415758 | ERR3415765 | ERR3415767 | ERR3415769 | ERR3415780 | ERR3415782 | ERR3415783 |
| ERR3415785 | ERR3415786 | ERR3415791 | ERR3435075 | ERR3435077 | ERR3435081 | ERR3435082 |
| ERR3435091 | ERR3435093 | ERR3435094 | ERR3435104 | ERR3435105 | ERR3435111 | ERR3435112 |
| ERR3435119 | ERR3435128 | ERR3525229 | ERR3525233 | ERR3525236 | ERR3525238 | ERR413154 |
| ERR413168 | ERR413176 | ERR413179 | ERR413196 | ERR413201 | ERR413203 | ERR413216 |
| ERR413218 | ERR413224 | ERR413227 | ERR413254 | ERR413254 | ERR413261 | ERR413265 |
| ERR413277 | ERR413286 | ERR413291 | | | | |

**Table S3: Oxidoreductase-associated Pfams among REBEAN identified enzymes**

| | Number of proteins | # of unique Pfams<br>All proteins | Enzymes | Oxidoreductases |
|---|---|---|---|---|
| SwissProt (Reference) | 539,105 | 14,953 | 5,183 | 798 |
| Identified putative oxidoreductases based read-level prediction | 275,290 | 6,021 | 3,338 | 663 |
| Novel putative oxidoreductases based read-level prediction | 32,030 | 2,942 | 1,672 | 381 |
| Novel putative oxidoreductases based read-level prediction and overall protein score | 4,901 | 1,043 | 647 | 204 |

### Table S4: Extreme environment samples used to evaluate REBEAN and comparator tools

| S. No | ENA SAMPLE ID | ENA RUN ID | Project | Pipeline version |
|---|---|---|---|---|
| 1 | ERS11432033 | ERR9456921 | Metagenomics insights into of Halophilic Communities inhabiting solar saltern in North Sinai, Egypt | 5 |
| 2 | ERS11432032 | ERR9456920 | Metagenomics insights into of Halophilic Communities inhabiting solar saltern in North Sinai, Egypt | 5 |
| 3 | SRS511419 | SRR1043601 | Saltern bacteroidetes | 4 |
| 4 | SRS511417 | SRR1043606 | Saltern bacteroidetes | 4 |
| 5 | SRS511471 | SRR1043669 | Saltern bacteroidetes | 4 |
| 6 | SRS511472 | SRR1043670 | Saltern bacteroidetes | 4 |
| 7 | SRS511470 | SRR1043668 | Saltern bacteroidetes | 4 |
| 8 | ERS1546719 | ERR2407577 | Metagenomics of Brava and Tebenquiche lakes | 4.1 |
| 9 | ERS3104179 | ERR3132464 | Diamante Lake Metagenome | 4.1 |
| 10 | ERS3040056 | ERR3083899 | Metagenomics of Ojo de Campo | 4.1 |
| 11 | ERS3104179 | ERR3132462 | Diamante Lake Metagenome | 4.1 |
| 12 | ERS3040055 | ERR3082345 | Metagenomics of Pozo Bravo microbial mats | 4.1 |
| 13 | ERS1546719 | ERR2407578 | Metagenomics of Brava and Tebenquiche lakes | 4.1 |
| 14 | ERS1546720 | ERR2437277 | Metagenomics of Brava and Tebenquiche lakes | 4.1 |
| 15 | ERS3036499 | ERR3079486 | Diamante Lake viral metagenome | 4.1 |
| 16 | ERS3104179 | ERR3132463 | Diamante Lake Metagenome | 4.1 |
| 17 | ERS3037840 | ERR3085921 | Yucra orchard metagenome | 4.1 |
| 18 | ERS3037841 | ERR3085930 | Severino orchard metagenome | 4.1 |
| 19 | SRS1379411 | SRR3341855 | Socompa Lake Stromatolite Shotgun Metagenomic | 4.1 |
| 20 | ERS2782947 | ERR2834522 | Kallisti Limnes subsea pools Metagenome Study | 4.1 |
| 21 | ERS2782947 | ERR2834520 | Kallisti Limnes subsea pools Metagenome Study | 4.1 |
| 22 | ERS2782948 | ERR2834314 | Kallisti Limnes subsea pools Metagenome Study | 4.1 |
| 23 | ERS2782947 | ERR2834521 | Kallisti Limnes subsea pools Metagenome Study | 4.1 |
| 24 | ERS2782948 | ERR2834315 | Kallisti Limnes subsea pools Metagenome Study | 4.1 |
| 25 | ERS2782948 | ERR2834316 | Kallisti Limnes subsea pools Metagenome Study | 4.1 |

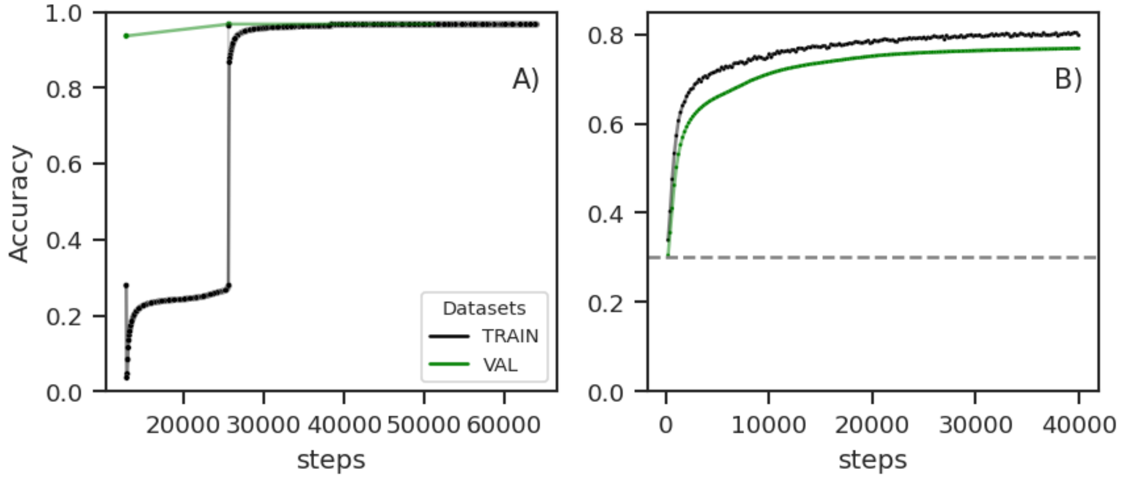


**Figure S1:** Training and validaiton accuracy of A) REMME and B) REBEAN. The gray line in plot B indicates the valdiation performance of REBEAN base model initiated with random weights instead of REMME weights.

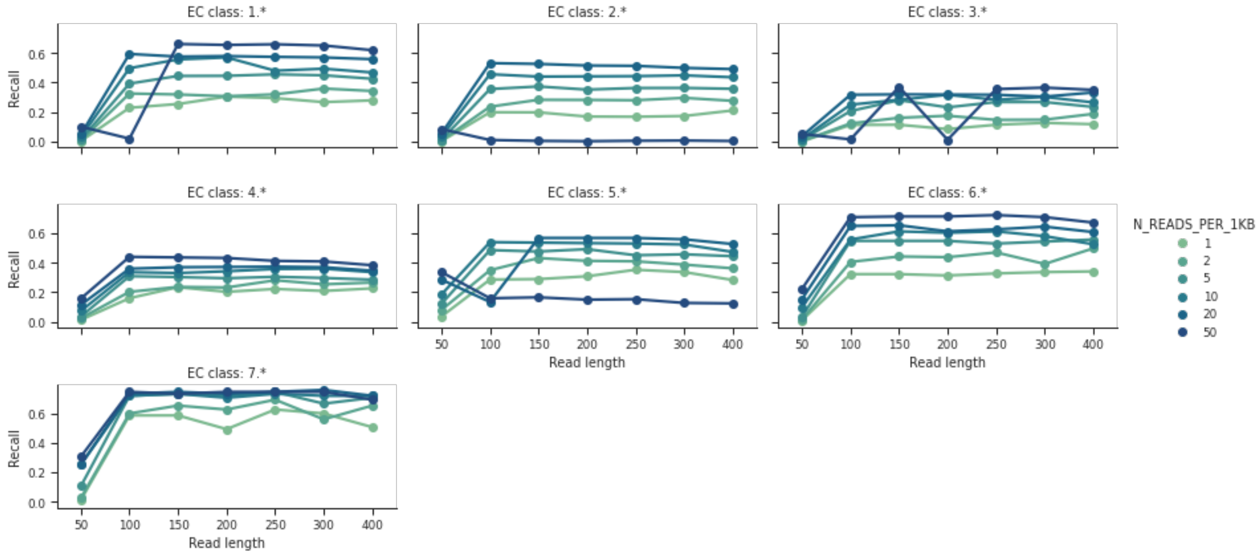


**Figure S2:** The effect of read length and read count on REBEAN's recall of known enzymes under each of seven EC First level classes.

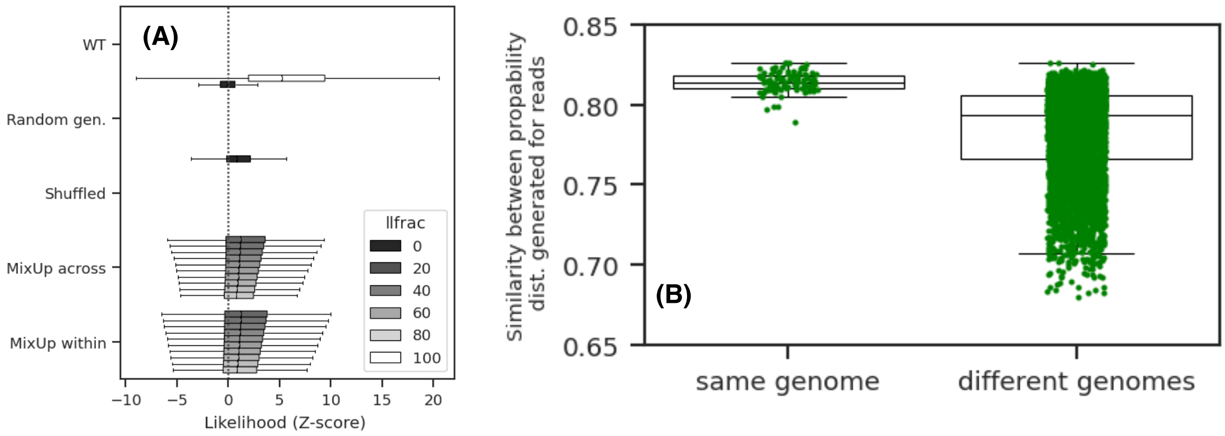


**Figure S3**: Scoring DNA sequences using REMME Decoder's probability matrix. We tested out REMME's interpretation of DNA sequences on read-sized fragments from 100 randomly selected GTDM genomes. We observed that the probability distribution generated by REMME decoder layers for a given read, uniquely encodes the given read in terms of emission probability which could be used to A) compute the likelihood of any other replacing read and B) to compute similarity between genomes.

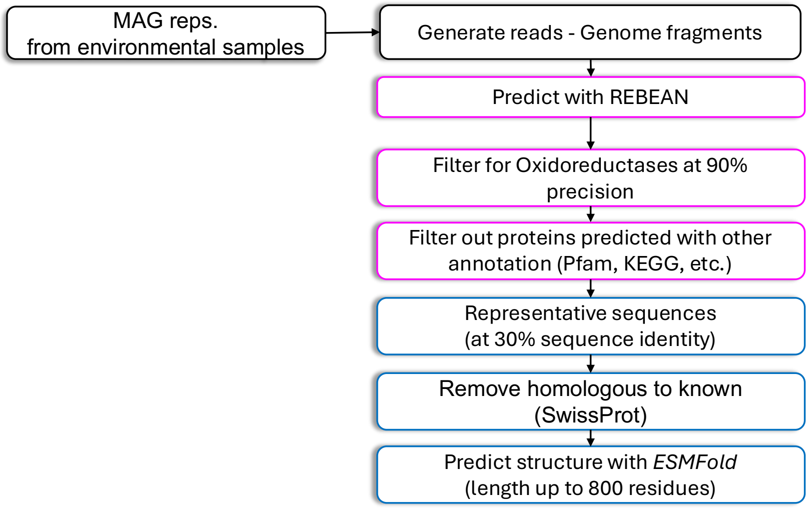


| **Reads** | **Proteins** | **Genomes** |
|---|---|---|
| 124 M | 11.3 M | 3,822 |
| 12.9 M | 1.13 M (10%) | 3,820 |
| 670, 637 | 407,428 (3.6%) | 3,820 |
| | 51,326 | |
| | 39,617 | |
| | **32,030** | |

**Figure S4:** Illustration of metagenomic enzyme mining using REBEAN-based pipeline

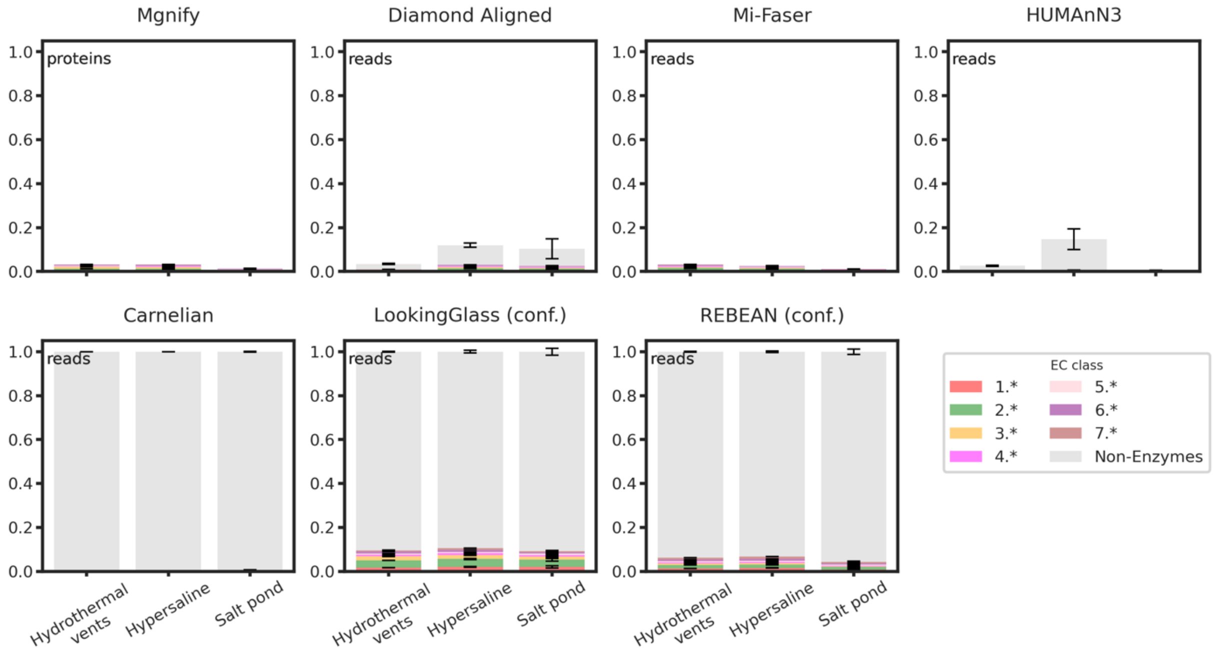


**Figure S5:** Fraction of reads annotated to first-level Enzyme Commission (EC) classes (1–7) or as non-enzymatic over 8 million metagenomic reads (**ExtremeMGset**), sampled randomly from three extreme environments. Annotation results are shown for REBEAN and other tools. For MGnify, the values represent the fraction of annotated proteins (out of ~500 million) from the same 18 metagenomic samples.

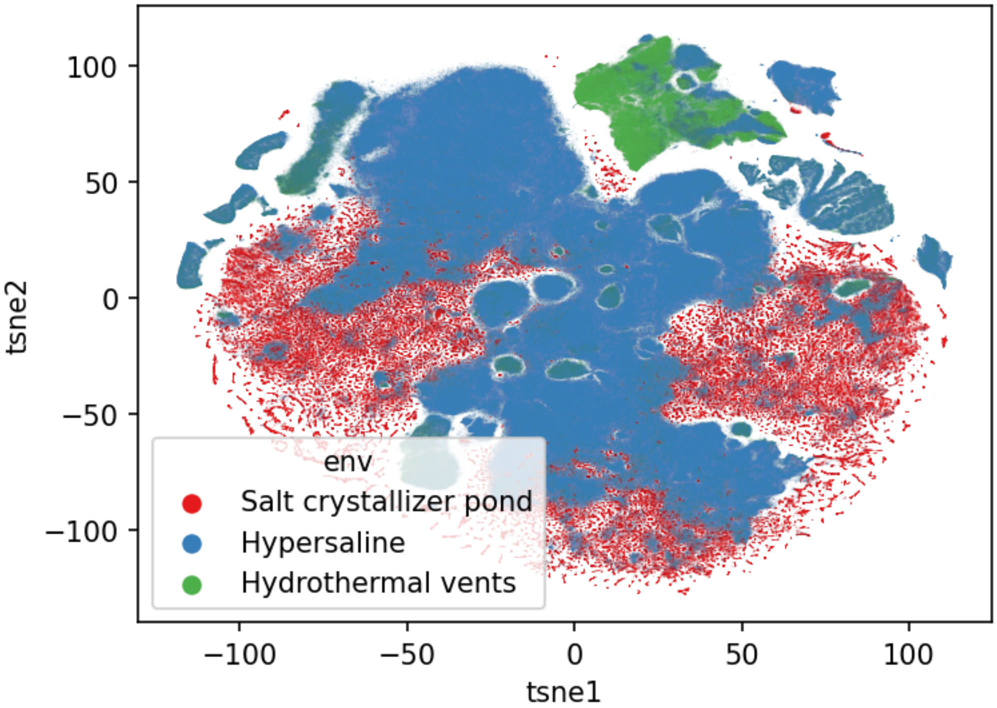


**Figure S6:** t-SNE projection of REMME embeddings for metagenomic samples from three extreme environments: Hydrothermal vents, Hypersaline and Salt crystallizer ponds.
